# Engineering “physically optimized” T cells for increased sampling of complex tumor microenvironments

**DOI:** 10.64898/2026.01.28.702394

**Authors:** Hongrong Zhang, Zhongming Chen, Guhan Qian, Christopher D. Zahm, Roberto Alonso-Matilla, Salena Fischer, Ingunn M. Stromnes, Beau R. Webber, Kevin W. Eliceiri, David J. Odde, Branden S. Moriarity, Paolo P. Provenzano

## Abstract

Pancreatic ductal adenocarcinoma (PDA) remains highly lethal, in part, because its dense fibroinflammatory stroma restricts therapy distribution, including adoptive T cell immunotherapies where direct interactions between T and carcinoma cells are essential for effective therapy. While T cell function must be maintained once effector-target engagement occurs, without inducing co-localization subsequent cytotoxic function steps cannot be undertaken. We therefore developed a strategy to “physically optimize” T cells to more effectively sample complex tumor volumes. Informed by pharmacologic perturbations and mathematical modeling we shifted T cell phenotype through expression of constitutively activated RhoA to increase cortical contractility, activation, migration, and sampling in PDA, while showing decreases in exhaustion markers. In CAR T cells this results in more efficient targeting through decreased sampling time and increased engagement with carcinoma cells, consistent with modeling predictions. This significantly increases T cell infiltration and distribution in PDA, resulting in improved tumor control in vivo, suggesting that this is an effective strategy to overcome stromal constraints, improve tumor engagement, and enhance the therapeutic performance of engineered T cell therapies in solid tumors.

## INTRODUCTION

Despite being the 14th most common cancer type, pancreatic ductal adenocarcinoma (PDA) is projected to become the second leading cause of cancer-related deaths due to its late-stage diagnosis, aggressive progression, and lack of effective therapeutic options (1–3). While a subset of localized PDA patients (∼20%) may benefit from surgical resection, the majority (∼80%) present with locally advanced or metastatic disease, making curative treatment unattainable. The current standard-of-care for these patients remains palliative chemotherapy. Although chemotherapy regimens offer a modest survival benefit, the overall prognosis remains extremely poor. Even among patients who undergo surgical resection followed by adjuvant chemotherapy, ∼80% experience disease relapse within five years (4, 5), underscoring the urgent need for more effective therapeutic strategies.

Immunotherapy, which harnesses the body’s immune system to target malignancies, is revolutionizing the treatment landscape for hematologic cancers, but successes in solid tumors, to date, are much more limited. Among various immunotherapeutic strategies, modulating agents or adoptive cell transfer for T cells is a major focus due to their potent cytotoxic capabilities. However, PDA remains largely refractory to immunotherapy (6–10), largely due to its desmoplastic stroma, which creates both physical and biochemical immunosuppressive barriers that restrict T cell distribution through the tumor volume and functional persistence (9, 11–14). Successful tumor elimination by cytotoxic T cells require a series of coordinated steps, including sufficient activation or engineering of effector T cells, infiltration into tumors, sampling of the entire tumor volume through heterogeneous tumor microenvironments (TMEs), and direct interaction with malignant cells while retaining sufficient function. However, the dense stroma in PDA—composed presents a formidable challenge to effective T cell migration, often leading to zones of immune exclusion (11, 15). Even in cases where T cells do infiltrate PDA, their distribution is restricted, resulting in many regions of the tumor that are free from anti-tumor immune dynamics (12, 16–18). As such, this spatial segregation between effector T cells and tumor cells represents a major mechanism of immune resistance in PDA. Indeed, while T cell function must be intact and maintained once effector-target engagement occurs, without successful effector-target co-localization the subsequent critical steps to kill transformed cells cannot be undertaken.

Developing strategies to enhance cytotoxic T cell migration through mechanically and chemically complex tumor landscapes is paramount for improving T cell immunotherapy efficacy in PDA. To start addressing this critical limitation we previously defined how perturbing the amoeboid-mesenchymal hybrid phenotype, which is utilized by T cells during migration through structurally complex environments, toward a more amoeboid state can improve T cell migration in PDA (11). In 3D environments and live tumors, increasing microtubule instability drives increased Rho pathway-dependent cortical contractility, resulting in a more ameboid shifted hybrid T cell phenotype that more effectively moves through and interrogates 3D matrix and tumor volumes (11). Hence, this work provides initial design criteria for engineering strategies to enhance T cells movement through desmoplastic solid tumors that can be part of an effective strategy to enhance efficacy of T cell therapeutics. However, although transient pharmacological activation of RhoA has been shown to increase T cell speed and overall motility in vitro, a sustained enhancement in migration and function is required to achieve persistent tumor sampling in vivo. Therefore, here we sought to test the hypothesis that cell engineering to harness RhoA signaling can enhance T cell motility to reduce immune exclusion. To test this hypothesis, here we generated T cells with high levels of constitutively activated Rho, RhoA(Q63L) and evaluated their phenotype and migration behaviors experimentally and through physics-based modeling. Functionally, RhoA(Q63L) T cells exhibit enhanced motility across multiple platforms, without affecting proliferation or viability. Importantly, RhoA(Q63L) T cells have a more activated effector phenotype while showing reduced levels of T cell exhaustion markers. In the therapeutic setting we find that integrating RhoA(Q63L) into chimeric antigen receptor (CAR) T cells results in increasing sampling and engagement with carcinoma cells, significantly higher CAR T cell numbers and distribution in PDA and greatly reduced tumor burden, defining a novel strategy to overcome physical barriers in PDA and improve immunotherapy outcomes.

## RESULTS

### T Cell Migration is Restricted in PDA Tumor Microenvironments

PDA is notoriously resistant to immunotherapy, in part, due to CD8+ T cell scarcity and profoundly immunosuppressive TMEs (12, 19, 20), with histological analyses consistently showing minimal to no T cell presence in many regions of PDA tumors (12, 16–18). However, histological analysis provides only a static snapshot of T cell distribution, offering little insight into the dynamic behavior of T cells within the TME. Thus, while prior studies have characterized the lack of T cell accumulation in many regions of PDA, they have not presented time course movement data to address whether T cells experience severe motility constraints that prevent effective sampling of the entire tumor volume to facilitate engagement with carcinoma cells. Therefore, to directly assess CD8+ T cell movement in real-time, we examined murine T cell motility in live PDA tumor slices from *Kras^LSL-G12D^;Trp53^LSL-R172H^;Pdx1-Cre;Rosa26^LSL-tdTomato/+^ (KPCT)* mice as we have described ((21); Fig. 1A) and compared it to migration in less complex, and much more migration permissive, 3D collagen matrices where T cells can effectively sample the matrix volume over time. T cell 4D (x,y,z,time) migration was captured using multiphoton-laser scanning microscopy (MPLSM) to simultaneously generate multiphoton excitation (MPE) of fluorescence and second harmonic generation (SHG) of fibrillar collagen. Within the live PDA tumor slice, both wild-type and therapeutic (engineered with a TCR against the PDA tumor-associate antigen mesothelin (22, 23)) T cell movement is highly restricted in many cases compared to T cells movement in more porous less dense 3D collagen matrices, where T cells exhibit significantly higher movement (Fig. 1B, Supp. Fig. 1 and Supp. Movies 1 and 2), consistent with robust physically and chemically immunosuppressive TME features in PDA. Even after 8 hours in the tumor slice, TCR-T cells migrated over significantly less than those in the collagen gel in just 1 hour, highlighting the profound limitations imposed by the PDA microenvironment. Indeed, quantitative analysis confirmed a major reduction in motility coefficient (Fig. 1C), total displacement (Fig. 1D), and track confinement ratio (Fig.1E) in tumor slices compared to the collagen matrices.

**Figure 1:**
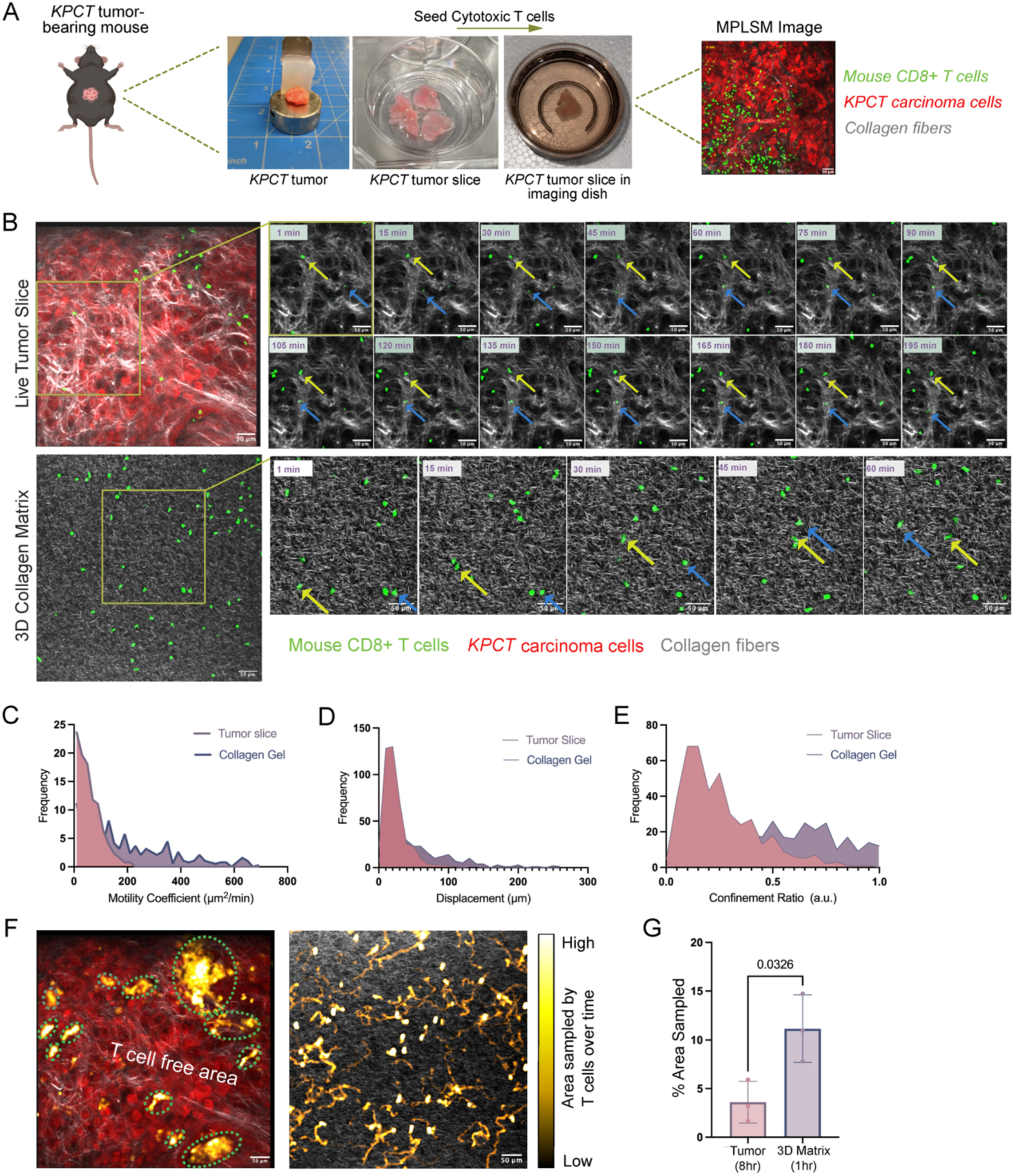
Limited T cell migration in PDA TMEs. **A**. Schematic of the live tumor workflow used to assess T cell migration in live mouse KPC tumor slices. **B**. Representative time-lapse images of CD8+ TCR-T cell migration within a live mouse PDA tumor slice over 8 hours (top) and within a 3D collagen matrix over 1 hour (bottom). Tracked cell positions are indicated by yellow and blue arrows. **C-E**. Distribution of T cell motility coefficient (C), displacement (D), and track confinement ratio (E) in tumor slice (pink) and 3D collagen gel (purple). **F.** Representative images showing spatial sampling by T cells in live mouse PDA tumor slice (8 hours) and in 3D collagen matrix (1 hour). Brighter color indicates higher T cell presence. **G**. Quantification of total area covered per field of view overtime by CD8+ T cells in tumor slice (Left, 8 hours) and in 3D collagen matrix (right, 1 hour).

To further quantify the extent of T cell restriction in the PDA tumor microenvironment, we analyzed the total area covered by T cells within the tumor slice compared to the collagen gel matrix. Over an 8-hour period, T cells in the tumor slice covered less than 5% of the total field of view (FOV), whereas T cells in the collagen gel covered >10% of the FOV in just 1 hour (Fig. 1F,G). In addition to reduced total area coverage, the distribution of T cells within the PDA tumor slices was also highly uneven, with large portions of the tumor remaining completely devoid of T cell sampling. In contrast, T cells in the collagen gel exhibited a more uniform distribution, indicating that they were able to freely migrate and explore their environment without encountering significant physical constraints (Fig. 1F,G and Supp Fig. 1). The presence of extensive T cell-free zones in the PDA tumor slices suggests that tumor-associated barriers not only restrict movement but may also create exclusion zones where T cells are unable to penetrate, further limiting effective immune surveillance within solid tumor environments.

### Engineering T cells with constitutively active Rho

We have previously demonstrated that pharmacological activation of RhoA significantly increases T cell speed and overall motility in 3D matrices (11). Based on these findings, we hypothesized that genetically enhancing RhoA activity in T cells would improve their migration within tumors, ultimately enhancing their ability to effectively sample and interact with carcinoma cells. The RhoA Q63L mutation, which replaces glutamine with leucine at position 63, results in a constitutively active form of RhoA by preventing GTP hydrolysis, thereby leading to constitutively activated RhoA (Fig 2A).

**Figure 2:**
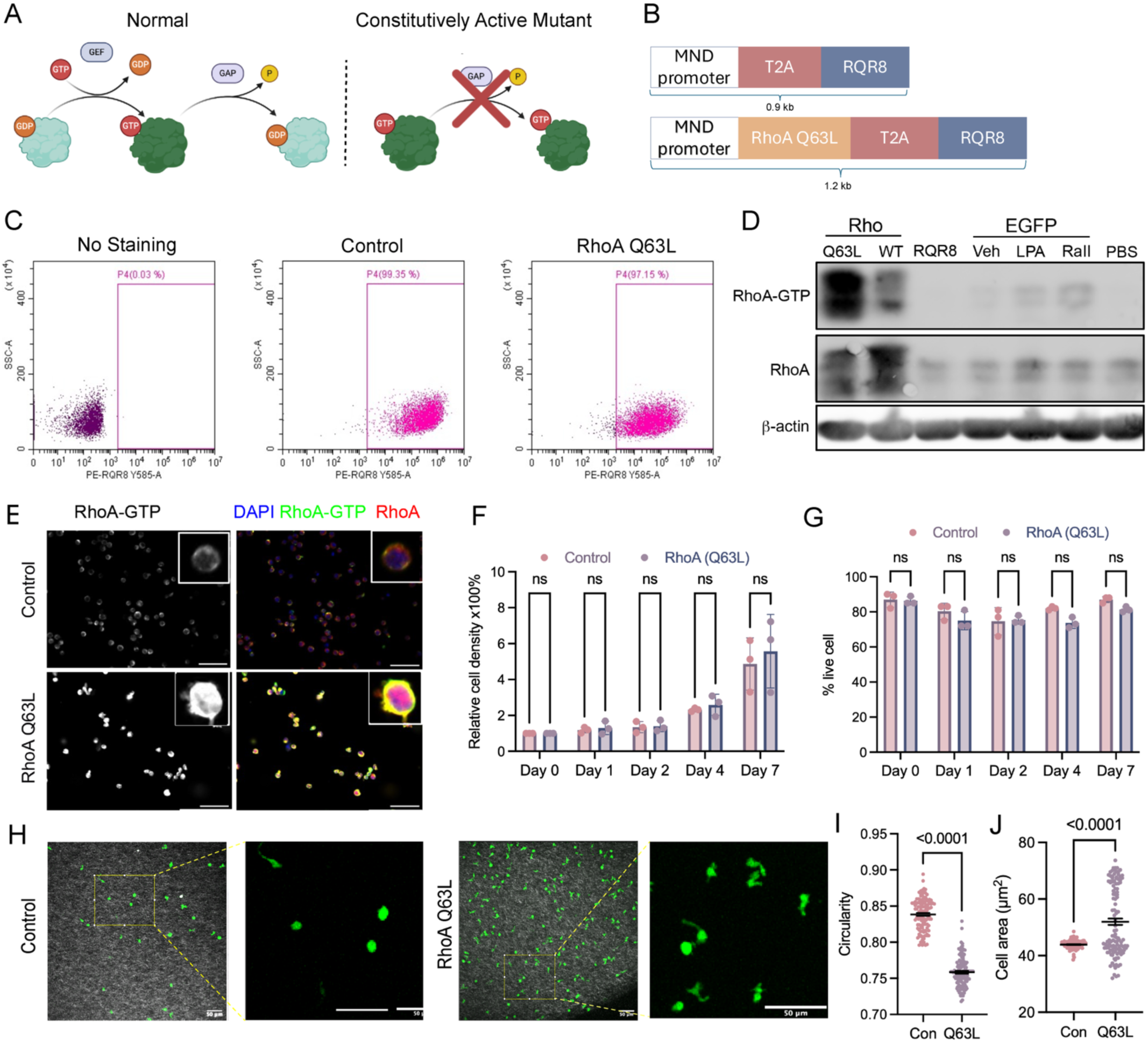
Generation and characterization of human RhoA(Q63L) T cells. **A.** Diagram illustrating the effect of RhoA(Q63L) mutation on GTPase regulation. **B.** Schematic of lentiviruses construct encoding control (top) and RhoA(Q63L) mutant (bottom), all driven by the MND promoter. T2A and P2A encode two self-cleavage peptides. RQR8 is used for detection and purification of transduced cells. **C.** Representative flow cytometry data of purified RQR8+ T cells using PE -conjugated anti-CD34 (QBend10) antibody. **D.** Representative Western blots of whole cell lysate and RhoA-GTP pulldown samples with RBD beads with antibodies against RhoA and β-actin. Rho activating treatments were included to assess basal protein expression and activation levels in T cells. **E.** Representative immunofluorescence images of T cells stained with RhoA (red) and RhoA-GTP (green) antibodies, and counter stained with DAPI (blue). Scale bar: 50μm. **F.** Proliferation rate and viability of human T cells expressing control or RhoA(Q63L) over 7 days. **G.** Representative images of infiltrated T cell morphology in 3D collagen gels. Quantification of cell morphology parameters demonstrated reduced circularity and increased cell size in RhoA(Q63L) T cells compared to the control. Each dot represents an individual cell pooled from three independent experiments. Data are presented as mean ± s.e.m.

To investigate the effects of RhoA(Q63L) mutant on T cell phenotype and migration, we transduced human T cells (mixed CD4+ and CD8+ populations) with a lentivirus encoding RhoA(Q63L) mutant and RQR8 under MND promoter, which initiates strong downstream gene expression in T cells (Fig. 2B). The RQR8 is an epitope-based marker that integrates CD34 and CD20, allowing positive selection of the successfully transduced cells, assuring high purity. To assess potential genetic variation, we prepared three batches of the RhoA Q63L mutant and control T cells from three donors. After purification, transduced T cell purity was assessed using flow cytometry, and show > 97% purity (Fig. 2C). Examination of RhoA-GTP levels in purified T cells with RhoA-GTP pulldown assay demonstrates high levels of active GTP bound Rho and total RhoA that are over 51 times and 11 times higher, respectively, than the RQR8 control (Fig. 2D). Furthermore, we note that basal levels of activated Rho are very low. Even under potent Rho stimulation conditions with LPA or RaII, we observe only modest increases in active Rho relative to activation from constitutively active Rho (Fig. 2D). To confirm this behavior, we further investigated the distribution of active RhoA using immunofluorescence. Consistent, with pull-down results, specific staining for GTP bound RhoA confirms that RhoA-GTP levels are strongly elevated Q63L expressing cells, with GTP bound Rho at higher levels near the cytoplasmic membrane compared to total RhoA protein that are more distributed throughout the cytoplasm (Fig. 2E). Overall, these results demonstrate highly pure populations of T cells expressing constitutively active Rho.

To ensure that the Q63L mutation does not negatively impact T cell proliferation or viability, we monitored T cell expansion in culture over 7 days (Fig. 2F,G). Quantification of cell number and viability showed no difference between the RhoA(Q63L) modified T cells and Control cells, indicating the Q63L mutation does not impair cell cycle progression or survival. However, RhoA Q63L modified T cells do display distinct morphological changes in 3D matrices. MPE+SHG imaging of the live T cells in 3D collagen matrices reveals that the Q63L modified T cells displayed a more elongated cell shape and increased cell size compared to the unmodified cells (Fig.2H), consistent with our previous findings that Rho activation shifts the amoeboid-mesenchymal phenotype balance that T cells employ while navigating fibrous matrices (11). Along these lines, our observations are consistent with a phenotype shift where T cells employ ameboid protrusion dynamics with cell elongation within the amoeboid-mesenchymal hybrid phenotype to enhance 3D migration through fibrous networks (11).

### Constitutively active Rho enhances T cell migration

Given the observed phenotype shift, we first sought to determine if constitutive Rho activation is altering cortical contractility that regulates the hybrid phenotype balance (11). To measure cortical contractility in live T cells we employed FLIPPER-TR technology with multiphoton fluorescence lifetime imaging microscopy (FLIM), where imaging demonstrates a significant increase cortical tension in RhoA(Q63L) expressing T cells (Fig. 3A,B), suggesting that Rho(Q63L) T cells may have increased capacity for 3D migration. To test this hypothesis and specifically evaluate the consequences of increasing cortical contractility on T cell migration we first performed mathematical modeling, using a recently developed physics-based model for hybrid T cells migration behaviors (24). Interestingly, model analysis predicts that increasing cortical contractility increases leading edge membrane elastic energy that drives amoeboid protrusion dynamics, results in increases in cell displacement, and overall migration speed (Fig. 3C)

**Figure 3:**
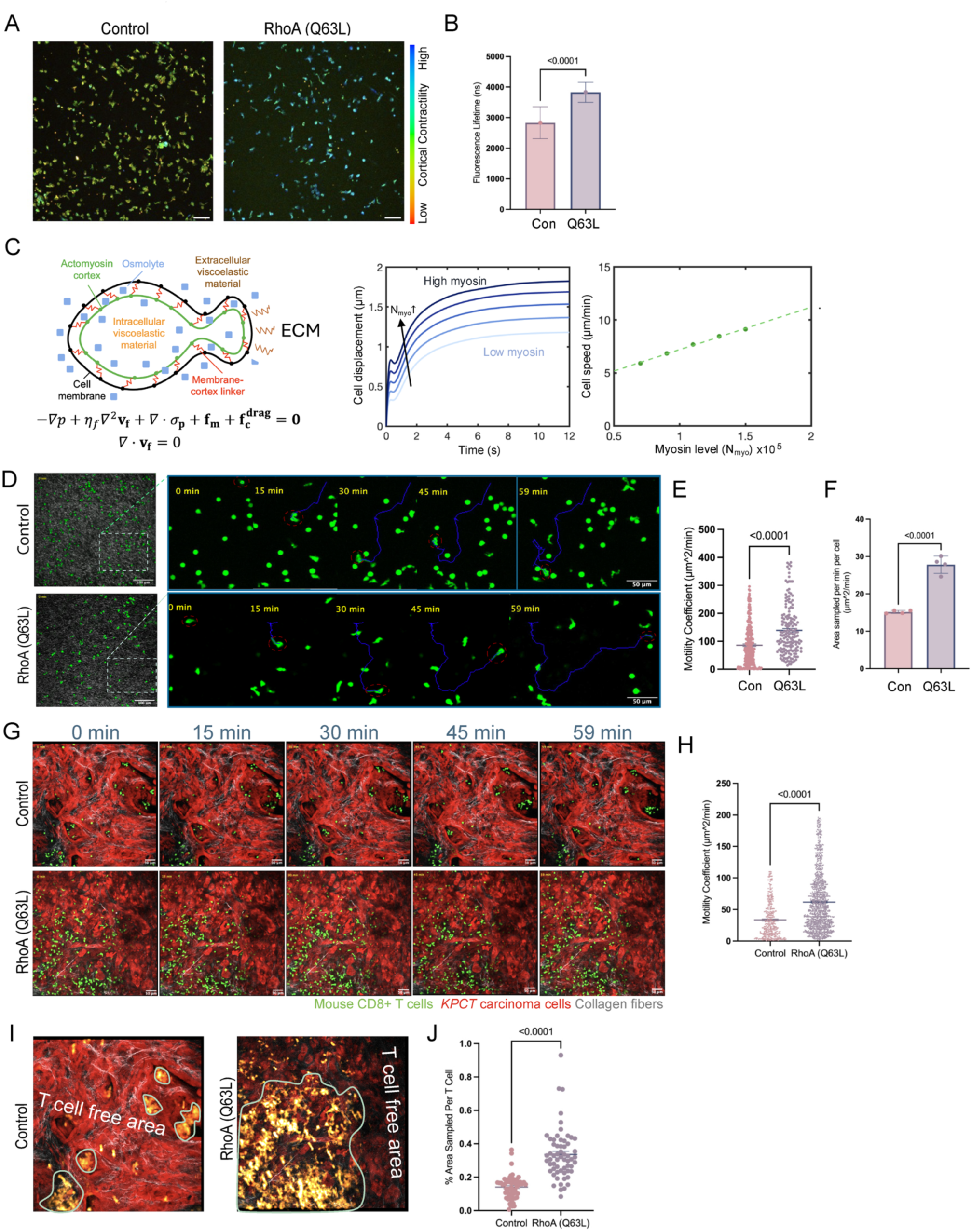
RhoA(Q63L) mutation significantly increases motility in both human and murine T cells. **A.** Fluorescence-lifetime images of control and RhoA(Q63L) T cells with Flipper-TR to assess cortical membrane tension. Cells are color-coded by fluorescence lifetime (red, shorter lifetimes; blue, longer lifetimes). **B.** Quantification of average fluorescence lifetimes across Z-stacks from Flipper-TR–stained T cells shows significantly longer lifetimes in RhoA(Q63L) cells, consistent with increased cortical contractility. **C.** Physics-based mathematical modeling predicts that increased cortical contractility results in increased T cell migration. **D.** Multiphoton time-lapse images of human control and RhoA(Q63L) T cells migrating in 3D collagen matrices. Representative single-cell migration tracks are highlighted in blue. Images were acquired every 1 min for 1 h. Colors: green, T cells; gray, collagen fibers. **E.** Motility coefficient for human control and RhoA(Q63L) T cells migrating in 3D collagen matrices. Each dot represents an individual tracked T cell. **F.** Average area sampled per frame by T cells migrating in 3D collagen matrices. n = 4 biological replicates. **G.** Multiphoton time-lapse images of murine control and RhoA(Q63L) and CD8⁺ T cells migrating within live *KPCT* pancreatic tumor slices. Colors: red, PDA carcinoma cells; green, T cells; gray, fibrillar collagen. **H.** Motility coefficients of murine control and RhoA(Q63L) CD8⁺ T cells migrating in tumor slices, showing a significant increase in motility in RhoA(Q63L) T cells. **I.** Representative multiphoton microscopy images showing the cumulative area sampled by infiltrating T cells in tumor slices over a 2-h period. Regions of high T-cell presence are outlined in green. **J.** Quantification of the percentage of area sampled per field of view per T cell. Data are presented as mean ± s.e.m. Statistical comparisons were performed using unpaired two-tailed t-tests with Welch’s correction.

To experimentally assess the impact of constitutively active RhoA (Q63L) on T cell motility, we first analyzed migration in a 3D collagen matrix system, which serves as a baseline assay for intrinsic migratory potential in the absence of chemotactic gradients (Fig. 3D). Human CD8⁺ T cells expressing RhoA(Q63L) exhibit significantly increased motility compared to control cells, as measured by an ∼2-fold increase in motility coefficient and a higher track confinement ratio (arbitrary units from 0 to 1, where 1 indicates a straight trajectory, i.e. more directional persistence; Fig. 3D,E and Supp. Fig 2). Importantly, RhoA(Q63L)-modified T cells also sample a larger area over time, consistent with enhanced intrinsic migration (Fig. 3F). Thus, combined, these data suggest that increasing cortical contractility in T cells via overexpression of constitutively activated Rho significantly enhanced movement and has the potential to improve sampling throughout tumor volumes.

To test the impact of activated Rho in T cells that are navigating native tumor landscapes we employed the *KPC* model with murine T cells engineered to express constitutively active Rho. Similar to human T cells, mouse CD8⁺ T cells transduced with RhoA(Q63L) display similar proliferation rates and viability compared to controls, exhibit a more elongated morphology and enlarged cell size, and motility in 3D matrices (Supp. Fig. 3). Therefore, to capture behaviors in more complex and heterogeneous TMEs, we analyzed T cell dynamics in live PDA slices derived from *KPC* and *KPCT* mice. Within live PDA tumor, RhoA(Q63L)-modified T cells showed significantly motility coefficients (Fig. 3G,H and Supp. Movies 3 and 4), as well as increased velocity and directional persistence, compared to controls. Furthermore, quantification of the area examined by the T cells over 1 hour demonstrates that on average each RhoA(Q63L)-modified T cells were able to sample over 0.33% of the FOV, whereases each control T cell covered less than 0.14%, resulting in multiple T-cell free regions (Figure 3I,J). This results in Rho(Q63L) T cells being being able to sample over 50% of the FOV, whereases the control T cells covered less than 10%, resulting in multiple T-cell free regions. These findings indicate that RhoA constitutive activation promotes T cell migration not only in engineered 3D matrices but also within the structurally dense and immunosuppressive PDA TMEs. More importantly, RhoA(Q63L) this increased motility results in more effective tissue sampling and spatial distribution, suggesting that these physically optimized T cells have an increased capacity to explore, and perhaps treat, tumor volumes.

### Physically optimized T cells have increased activation and decreased exhaustion

Since T cells function depends on its activation state and metabolic state is a readout of T cell activation and effector function (25, 26), we first investigated whether RhoA(Q63L) alters T cell metabolism. Using multiphoton fluorescence lifetime imaging microscopy (FLIM), we measured nicotinamide adenine dinucleotide NAD(P)H fluorescence lifetimes (Fig. 4A). where FLIM of the endogenous metabolic co-enzyme reduced nicotinamide adenine dinucleotide (NAD(P)H) can capture the fraction of free and protein-bound co-enzyme to provide quantitative endpoints of T cell metabolism. Consistent with a previous report (25), activation with aCD3/aCD28 results in increased percent free NAD(P)H (α_1_), (Fig. 4A,B). Analysis of the Rho(Q63L) and unmodified T cells with and without aCD3/aCD28 T cell activation likewise reveals that the physically optimized T cells display a higher α_1_ percentage (Fig. 4A,B), indicating a shift in cellular metabolism toward a state associated with T cell activation and proliferation. Furthermore, aCD3/aCD28 activated RhoA(Q63L) T cells have the highest percentage of NAD(P)H α_1_, follow by Rho(Q63L) modified T cell without aCD3/aCD28 T cell activator antibody (Fig. 4B), suggesting that the addition of constitutively activated Rho promotes an activated T cell state.

**Figure 4:**
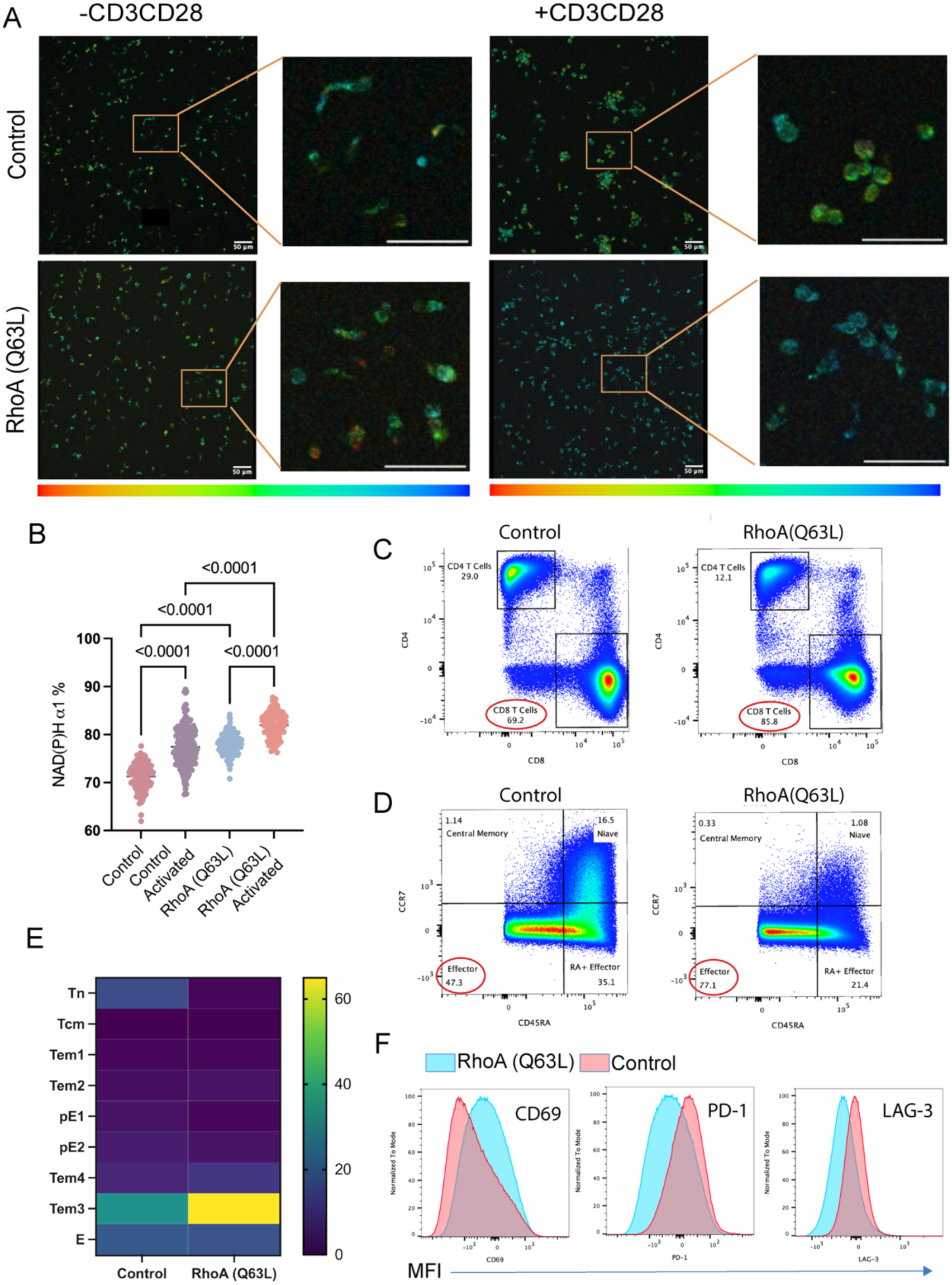
Effect of RhoA(Q63L) mutation on human T cell metabolism, differentiation, and phenotype. **A.** Fluorescence lifetime imaging of control and RhoA(Q63L) T cells inside 3D collagen matrices. Left: unstimulated; right: stimulated with anti-CD3/CD28 antibodies; Color bar indicates percentage of NAD(P)H α1from FLIM analysis (red: low percentage; blue: high percentage). **B.** Quantification of NAD(P)H α1 percentage in unstimulated and CD3/CD28-stimulated control and RhoA(Q63L) T cells. RhoA(Q63L) T cells showed significantly elevated baseline α1 levels, indicating increased activation. **C-D.** Multiparameter flow cytometry analysis of T cell subpopulations. RhoA(Q63L) modified T cells display a higher frequency of CD8⁺ T cells (C) and effector subsets (D) compared to control T cells. **E.** Heatmap of T cell percentage across differentiation stages: naïve (Tn), central memory (Tcm), effector memory (Tem), and terminally differentiated effector (E) cells, showing as heatmap. RhoA(Q63L) modified T cells exhibited a more differentiated phenotype, with the majority residing in the Tem3 (∼60%) and E (∼15%) subsets. **F.** Expression of exhaustion markers PD-1 and LAG-3 on T cells. RhoA(Q63L) modified T cells exhibit reduced PD-1 and LAG-3 expression compared to control T cells, suggesting a less exhausted phenotype.

To further investigate the effects of constitutively activated Rho on T cell function, we analyzed T cell activation, differentiation, and exhaustion status using multiparameter flow cytometry. Compared to unmodified controls, human RhoA(Q63L) T cells exhibit a higher percentage of CD8⁺ T cells and effector T cell subsets, suggesting that these cells are better at maintaining their cytotoxic and effector functions (Fig. 4C). Given this shift toward an enhanced effector phenotype, we next sought to determine whether expression of RhoA(Q63L) alters the differentiation stages of CD8⁺ T cells. CD8⁺ T cell differentiation can be ordered from naïve (Tn) cells to central memory (Tcm), effector memory (Tem), and terminally differentiated effector (E) cells, with each stage characterized by distinct cell surface marker expression. Using established gating strategies based on CD45RA, CCR7, CD27, and CD28 expression (27), we found that Q63L-modified T cells exhibit more differentiated subsets compared to the unmodified controls (Fig. 4D,E). The majority of Q63L-expressing T cells were found in the Tem3 (60%) and E (15%) stages, whereas unmodified T cells had a significantly lower percentage of Tem3 cells (<40%) and effector cells (<20%) (Fig. 4D,E). Additionally, the control group comprised a higher percentage of naïve T cells. Taken together, overexpression of constitutively active Rho in T cells promotes a shift toward functionally mature and specialized effector subsets, where highly differentiated T cells, particularly Tem3 and E subsets, are often associated with increased cytotoxic phenotype.

Importantly, analysis of expression levels for the T cell exhaustion markers PD-1 and LAG-3 revealed that RhoA(Q63L) T cells do not exhibit signs of exhaustion (Fig. 4F). Instead, these cells maintain a functional, cytotoxic phenotype with strong effector potential, distinguishing them from exhausted T cell populations typically observed in TMEs. These results suggest that, in addition to enhancing T cell migration, the Q63L modification drives differentiation toward a more functionally potent effector state while preventing exhaustion-associated dysfunction. This unique combination of enhanced mobility, differentiation, and resistance to exhaustion may provide a therapeutic advantage in improving the persistence and effectiveness of engineered T cells in solid tumors, and in particular PDA.

### Physically optimized CAR T cells display increased T cell – carcinoma cell interaction events

Having established that physically optimizing T cells through overexpression of constitutively active Rho enhances T cell activation and motility through complex TMEs, we sought to define the benefit in therapeutic CAR T cells. Thus, we designed and incorporated the mesothelin-CAR into human CD4+/CD8+ T cells (mesoCAR), with and without the RhoA(Q63L) (Supp. Fig. 4A), as mesothelin is an established tumor-associated antigen for PDA (23, 28). RhoA-GTP protein pulldown assays confirmed that the mesoCAR RhoA(Q63L) T cells have increased active Rho (Supp. Fig. 4B,C). Consistent with our previous findings, time-lapse imaging of the mesoCAR T cells in 3D collagen matrices show enhanced migration capacity with an ∼2-fold increase in motility coefficient (Fig 5A), as well as increases in speed, displacement, total distance traveled, and track confinement ratio (Supp. Fig. 5A-C).

**Figure 5:**
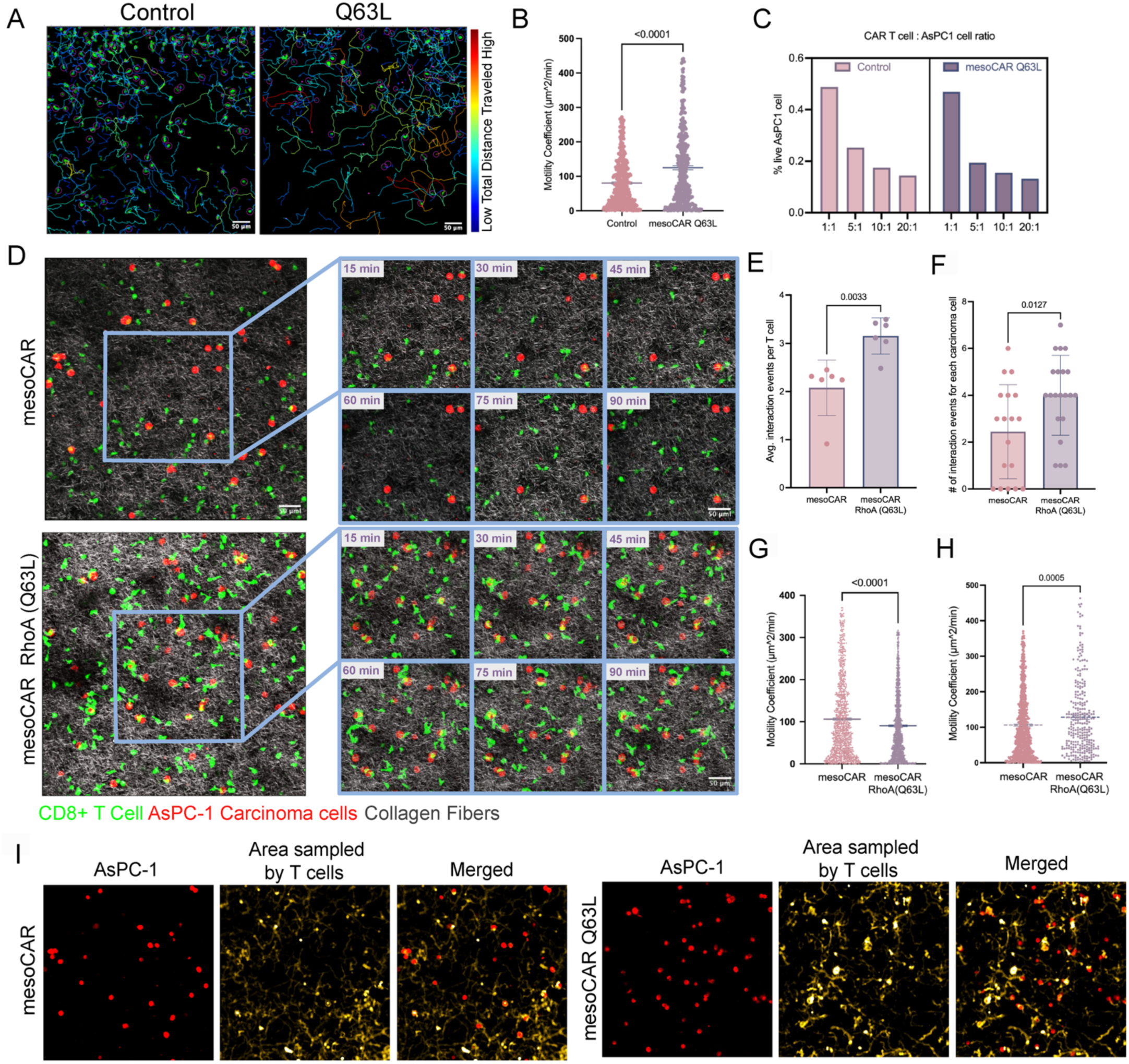
Incorporation of the RhoA(Q63L) enhances CAR T cell motility and carcinoma engagement. **A.** Migration of RhoA(Q63L) modified and control mesoCAR T cells in 3D collagen matrices. Migration trajectories are overlaid on time-lapse images and color-coded by total distance traveled (blue, short; red, long). RhoA(Q63L) modified CAR T cells display increased motility and more persistent movement. **B.** Quantification of mesoCAR T-cell motility coefficients in 3D collagen matrices. RhoA(Q63L) modified CAR T cells exhibit significantly higher motility coefficients than control CAR T cells. **C.** Quantification of 2D cytotoxicity. mesoCAR and mesoCAR RhoA(Q63L) modified T cells. Both CAR T-cell groups show similar increases in target-cell killing with increasing effector ratios. **D.** Multiphoton images of mesoCAR and mesoCAR RhoA(Q63L) modified T cells migrating and interacting with AsPC-1 cells in 3D collagen matrices. Colors: green, T cells; red, AsPC-1 cells; gray, collagen fibers. **E.** Quantification of T-cell–AsPC-1 interaction events per T cell for mesoCAR, and mesoCAR RhoA(Q63L) modified T-cell groups in 3D collagen matrices. **F.** Quantification of T-cell–AsPC-1 interactions events per carcinoma cell for mesoCAR and mesoCAR RhoA(Q63L) modified T-cell groups in 3D collagen matrices. **G.** Quantification of mesoCAR and mesoCAR RhoA(Q63L) modified T-cell motility in collagen matrices containing AsPC-1 cells. **H.** Quantification of mesoCAR and mesoCAR RhoA(Q63L) modified T-cell motility in collagen regions lacking AsPC-1 cells. **I.** Multiphoton microscopy images showing the cumulative area explored by mesoCAR, and mesoCAR RhoA(Q63L) modified T cells in AsPC-1–embedded 3D collagen matrices. Overlaid tracks and projected cumulative areas over 90 min are shown (red, AsPC-1 cells; yellow, T-cell area coverage). Data are presented as mean ± s.e.m. Statistical comparisons were performed using unpaired two-tailed t-tests with Welch’s correction.

To investigate the direct cytotoxicity of the mesoCAR +/− RhoA(Q63L) T cells we examined carcinoma cell killing in 2D culture conditions that facilitate close proximity of the CAR T cells with their target cancer cells, i.e. lack 3D barriers to effective T cell – carcinoma cell interactions. We performed these 2D killing of the AsPC1 PDA cancer cell line with E:T ratios of 1:1, 5:1, 10:1, and 20:1, where carcinoma cells cultured without T cells were used as a baseline and the percent survival for each of the coculture groups to the control was calculated. Our data show similar killing efficiency between the mesoCAR T cells +/− the Q63L modification (Fig. 5C and Supp. Fig. 6). Both groups displayed a trend in increased cancer cell killing as the ratio of T cells to carcinoma cells increases, suggesting the Q63L modification does not impair direct CAR-mediated cytotoxicity when cells are easily engaged. Thus, we next sought to evaluate CAR T – carcinoma cell interactions in 3D environments.

We embedded AsPC1 cells in collagen matrices, allowed the CAR T cells to infiltrate the matrices, and performed time-lapse imaging for 2 hours (Fig. 5D and Supp. Movie 5 and 6). T cell migration and morphology changes were tracked in 4D and we quantified T cell – carcinoma cell interaction events per T cell. Notably, on average the mesoCAR control group had 2 interaction events per T cell, whereas the mesoCAR Q63L modified group had 4 carcinoma cell interaction events per T cells, a two-fold increase (Fig. 5E). Likewise, analysis of the number of times a carcinoma cell was encountered by a T cells shows ∼2.4 vs. ∼4 for mesoCAR and mesoCAR Q63L groups, respectively (Fig 5F). Consistent with this behavior the motility coefficient of mesoCAR RhoA(Q63L) T cells is lower than that of the mesoCAR control T cells when carcinoma cells are present (Fig. 5G), in contrast to the superior motility in 3D matrices without AsPC1 cells (Fig. 5A,B) or in regions aways from AsPC1 cells (Fig. 5H), suggesting that the reduction in overall motility is due to increases in T-carcinoma cell interactions where T cells dwell instead of sampling. Thus, we conclude that when not interacting with carcinoma cells, Q63L-expressing T cells migrate faster, enabling them to locate targets more effectively. This faster search time results in a higher frequency of interactions, ultimately leading to a decrease in net motility due to engagement time with carcinoma cells. Indeed, this is consistent with theoretical predictions from Smoluchowski limit modeling parameterized from our 4D dataset that predicts mesoCAR T cells spend ∼88% of their time searching for carcinoma cells and ∼12% in contact (Supp. Modeling Materials). In contrast, mesoCAR Rho(Q63L) only ∼66% of their times searching and 34% of their time interacting the carcinoma cells. This is also supported experimental as analysis of the areas covered by T cells over the entire imaging period shows that mesoCAR Rho(Q63L) T cells are more concentrated and dwell around carcinoma cells compared to the control group (Fig. 5I). Combined this data suggests that increased motility of mesoCAR Rho (Q63L) T cells results in increased interactions between T cells and the target carcinoma cells.

### Physically optimized demonstrate improved therapeutic impact against PDA

To evaluate the therapeutic efficacy of the human RhoA(Q63L) engineered CAR T cells, we established an orthotopic pancreatic tumor model by grafting 1:2 ratio of AsPC1 cancer cells with human CAFs in 3 mg/mL collagen into the pancreas of NSG mice (Fig. 6A). Mice were enrolled for intervention studies when the tumor reached 5-6 mm in the longest dimension. Tumor growth was monitored by MRI (Fig 6A,B). Prior to treatment tumors volume remained comparable across all groups, confirming equivalent baseline tumor burden before treatment randomization (Fig. 6C). At the start of treatment mice received intraperitoneal injections of either mesoCAR T cells, mesoCAR RhoA(Q63L) T cells, or no T cell control. Tumors in the no T cell control group exhibited rapid growth (Fig. 6C,D). Treatment with the mesoCAR T cells showed a trend of reduced tumor volume. In contrast, treatment with mesoCAR RhoA(Q63L) T cells demonstrates a significantly reduced tumor growth (Fig. 6D)., indicating that the RhoA(Q63L) modification enhances the therapeutic efficacy of mesoCAR T cells in an orthotopic PDA model.

**Figure 6:**
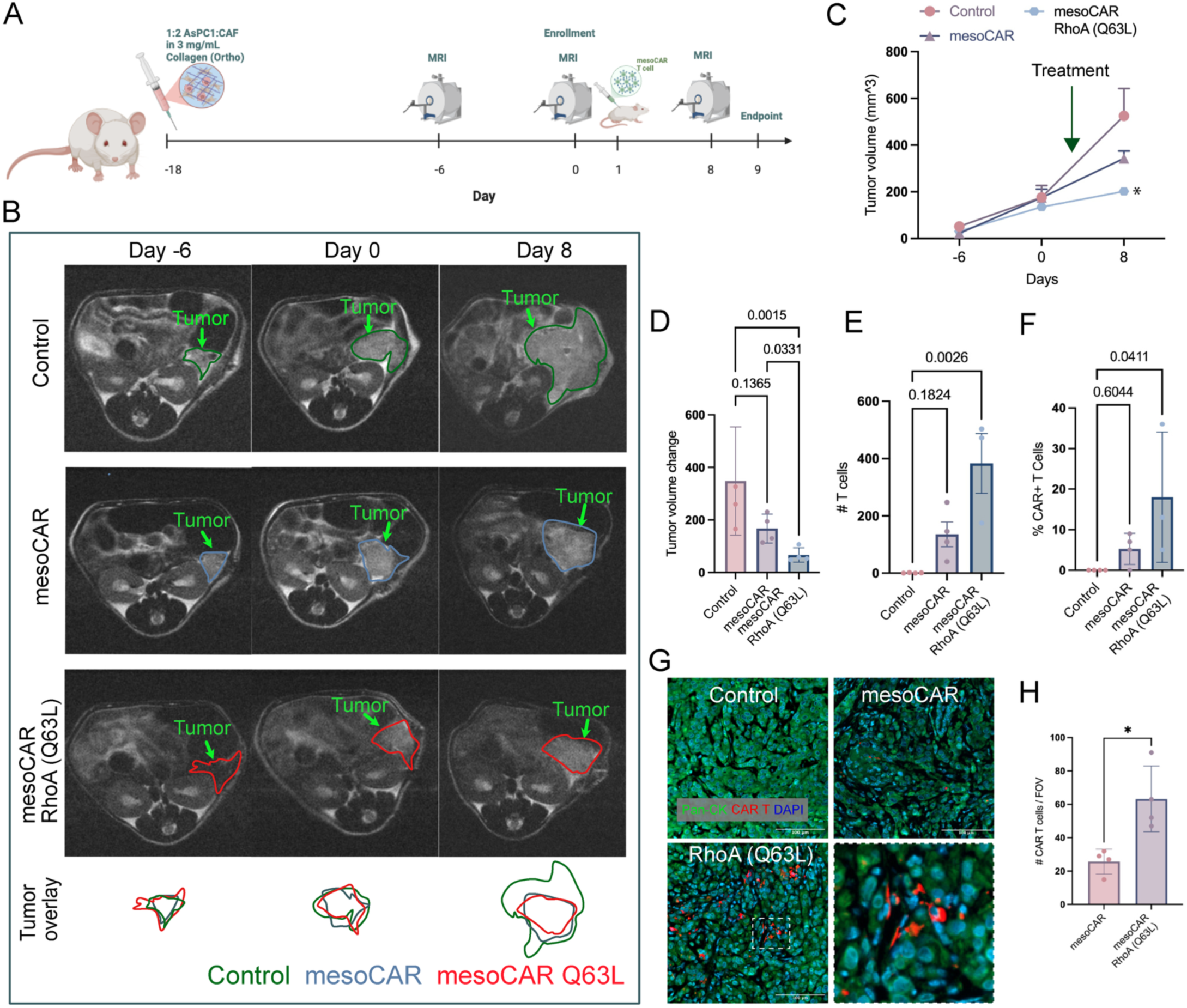
Q63L modification enhances CAR T-cell infiltration and tumor suppression. **A.** Schematic of the experimental design. AsPC-1 cancer cells and human CAFs (1:2 ratio) embedded in 3 mg/mL collagen gels were orthotopically implanted into the pancreas of NSG mice (Day –18). Tumor progression was monitored by MRI on Days –6, 0, and 8. On Day 1, mice (n = 4 per group) received intraperitoneal injections of mesoCAR T cells, mesoCAR Q63L modified T cells, or no–T-cell vehicle control. **B.** Representative MRI images of tumors at Days –6, 0, and 8 for each treatment group. Tumors were outlined and color-coded by group: green, control; blue, mesoCAR; red, mesoCAR Q63L modified. **C.** Tumor-volume trajectories across time. **D.** Quantification of changes in tumor volume after treatment. **E–F.** Flow-cytometry analysis of digested tumors showing the number (E) and percentage (F) of CAR T cells among all live cells. The mesoCAR Q63L modified group displays a significantly higher number of infiltrating CAR T cells relative to mesoCAR. **G.** Representative immunofluorescence images of FFPE tumor sections stained for pan-cytokeratin (green; tumor), RQR8 (red; CAR T cells), and DAPI (blue; nuclei). mesoCAR Q63L modified T cell–treated tumors show increased intratumoral CAR T-cell presence. Scale bar, 100 μm. **H.** Quantification of CAR T cell numbers per field of view in tumor immunofluorescence sections. Data are presented as mean ± s.e.m. Statistical significance was determined using one-way ANOVA with Tukey’s post hoc test. For all pairwise comparisons, unpaired two-tailed t-tests with Welch’s correction were used.

To access whether the improved tumor control is related to increased T cell infiltration, we first quantified intratumor CAR T cells by flow cytometry. Analysis of digested tumor tissues to create single cell suspensions revealed a robust and significant increase in the absolute number and percentage of CAR T cells, using the RQR8 marker, among all live cells in the mesoCAR Q63L group compared to the mesoCAR group (Fig. 6E,F). Consistent with these findings, immunofluorescence analysis of FFPE tumor sections stained for pan-cytokeratin (tumor), RQR8 (CAR T cells), and DAPI (nuclei) show greater accumulation of CAR T cells in PDA for mesoCAR RhoA(Q63L) treated mice relative to the mesoCAR control group (Fig. G,H). Together, these results demonstrate that incorporation of the RhoA(Q63L) modification enhances infiltration and distribution of CAR T cells and increases antitumor efficacy in vivo, consistent with our in vitro findings of improved T cell motility and tumor sampling.

## DISCUSSION

Our study demonstrates that overexpression of constitutively activated RhoA through the Q63L mutation reprograms T cells to overcome physical and molecular constraints in PDA by enhancing their motility and tumor sampling capacity without compromising viability or effector potential. We provide direct evidence that the PDA TME imposes profound biophysical barriers on T cell migration. While unmodified T cells exhibited severely limited displacement and spatial confinement, consistent with prior reports of immune exclusion and elevated interstitial pressures in PDA (12, 16–18, 29), RhoA(Q63L) T cells display significantly improved migration and therapeutic impact. These data support the hypothesis that targeting cell-intrinsic migratory programs can bypass aspects of stromal immune exclusion, a major limitation in current T cell-based approaches for solid tumors. Mechanistically, constitutive RhoA activation promotes a transition toward a more amoeboid migration mode for elongated cells, characterized by elevated cortical tension, suggesting that alternative approaches to modulate contracilitiy and phenotype may provide additional avenues to develop next-generation “physically optimized” T cells. Moreoever, while this is likely not limited solely to Rho signaling, defining the signaling alerations associated with constitutively activating Rho is warranted. Overall, it is clear that employing cell engineering approaches to enhance the physical function of therapeutic cells has the potential to profoundly improve immune therapy efficacy in solid tumors.

Our results also align with a broader body of literature advocating for stromal reengineering to improve therapeutic efficacy in PDA. While several stroma-targeting therapies such as enzymatic degradation of hyaluronan, inhibition of Hedgehog signaling, and suppression of collagen synthesis have largely focused on increasing chemotherapy distribution and altering intrinisic immune dynamics, studies to rigorously define how STT can improve adoptive cell transfer therapy are needed. Certainly, if cells can be engineered to overcome stromal barriers then reducing the barriers will be likewise beneficial. While it is possible that cell engineering approaches to physically optimize immune therapeutic cells, or perhaps cocktails of distinct physically optimized cells to overcome heterogenous barriers, may be sufficient in some cases, bypassing the need to deplete or remodel stromal components, it appears likely that rational combinations of stroma normalizing STT and physically optimized cells will be needed for broad curative impact of both primary and metastatic disease, particularly since TMEs can evolve, highlighting the critical need to define spatiotemporal dynamics of immune therapies in solid tumors. Along these lines, while a critical step, we acknowledge that improved migration alone, while critical, is unlikely to fully overcome the complex barriers to immunotherapy efficacy in PDA. While the physical optimization here with RhoA(Q63L) addresses a major physical bottleneck by enhancing T cell distribution within PDA, additional strategies will likely be necessary to achieve robust and durable antitumor responses. Combining physical optimization with approaches such as reinforcing T cell proliferation, survival and/or killing potency, resistance to exhaustion, or ability to modulate the immunosuppressive milieu (e.g. altering checkpoint blockades, metabolic reprogramming, or depletion of immunosuppressive myeloid cells) are likely required to fully unlock the therapeutic potential of T cells for solid tumor treatment, and particularly for robustly fibrotic and immunosuppressive tumors like PDA.

In summary, our findings define a cell engineering approach to physically optimize T cells to overcome critical physical barriers in PDA, restoring their ability to migrate, survey, and engage tumor cells in densely fibrotic environments. This strategy may represent a broadly applicable next-generation platform to enhance adoptive T cell therapies for solid tumors characterized by stromal exclusion and immune suppression. Moving forward, a comprehensive strategy that integrates enhanced migration with reinforcement of effector function and modulation of the suppressive TME is key to unlocking the full potential of immunotherapy in solid tumors and currently intractable malignancies such as PDA.

## MATERIALS and METHODS

### Human T cell culture

Human CD4+ and CD8+ T cells were isolated from healthy human donors and stimulated with anti-human CD3/CD28 beads in Optimizer human T cell expansion medium supplemented with 30 U/mL of human IL-2, IL-7, and IL-15, and transduced with lentivirus at MOI=20 on day 2. The transduced T cells were purified on day 4 with PE-CD34 QBEND/10 antibody (Thermofisher, Carlsbad, CA) and EasySep PE-positive selection kit (STEMCELL, Cambridge, MA). Cell purity was examined with flow cytometry thereafter. High purity T cells (purity>95%) were transferred to 24-well G-REX for cell expansion for additional 3 days.

### Mouse T cell isolation, activation, and culture

Mouse cytotoxic CD8^+^ T lymphocytes were extracted from C57/Bl6 wild-type or TRex genetically engineered mice (22) spleens using the EasySep CD8^+^ T Cell Isolation Kit (STEMCELLTM Technologies Inc., USA), following the manufacturer’s recommendation. Isolated CD8^+^ T cells are cultured and activated using the Dynabeads Mouse T-Activator CD3/CD28 (ThermoFisher Scientific, MA), following the manufacturer’s recommendation. If frozen T cell were used, the cells are thawed in a 37 °C water bath for 2 to 3 minutes and washed in 10 mL fresh pre-warmed Immunocult media to remove excessive DMSO in the medium. Thawed T cells are allowed to recover in fresh Immunocult medium supplemented with 30 U/ml IL-2 (STEMCELLTM Technologies Inc., USA) for at least 24 hours before subsequent experiments. All animal and cell work were approved by the University of Minnesota Institutional Biosafety Committee and followed institutional and NIH guidelines.

### Generation of human and mouse RhoA(Q63L) mutant T cells

Three lentiviral constructs were designed for the current studies. To prepare lentivirus encoding human RhoA(Q63L) and RQR8 for purification, the human amino acid codon optimized sequences of RhoA(Q63L) and RQR8 were linked with a T2A sequence and inserted to a pLV lentivirus backbone and driven by a synthetic viral promoter, MND (**M**yeloproliferative sarcoma virus enhancer, **N**egative control region-deleted, **d**l587rev primer-binding site substituted promoter). Vectors encoding overexpression RhoA-T2A-RQR8, and T2A-RQR8 were used as controls. For generating the mouse RhoA(Q63L) mutant T cells, cells were transduced with retrovirus constructs similar to the human constructs. Briefly, mouse amino acid codon optimized sequences of RhoA(Q63L) and RQR8 were linked with a T2A sequence and driven by the murine stem cell virus (MSCV) promoter. Both human and mouse RhoA(Q63L) and control vectors and the respective lentiviruses and retroviruses were prepared by VectorBuilder Inc (Chicago, IL).

### RhoA-GTP pulldown and Western blots

To examine RhoA-GTP induced by RhoA Q63L constitutional mutant, T cells were harvested for RhoA-GTP pulldown assay with RBD beads following manufacturer’s instructions (Cytoskeleton Inc, Denver, CO). RhoA-GTP and total RhoA were examined by Western blots with rabbit anti-human RhoA (1:500, Cell Signaling, Danvers, MA) and mouse anti-β-actin (1:1000, GeneScript, Piscataway NJ) antibodies, followed by secondary antibodies of IR680-conjugated donkey anti-rabbit IgG (1:2500) and IR800-conjugated donkey anti-mouse IgG (1:2500, LI-COR, Nebraska, USA).

### Immunofluorescence

T cells were fixed with 1% paraformaldehyde for 10 minutes and washed with PBS once. The fixed T cells were resuspended on deionized water and transferred to positively charged glass slides. The slides were air dried in a running cell culture hood, typically, in 10 to 20 minutes, to attach T cells on slides. The T cells on the slides were subsequently permeabilized with 0.3% Triton X-100 in PBS for 10minutes, and blocked with 1% FBS, 0.5% BSA for 60 minutes in room temperature. To examine total RhoA and RhoA-GTP, 1:50 rabbit anti-total RhoA (Novus Bio,Centennial, CO) and 1:100 mouse anti-RhoA-GTP (NewEast Biosciences, King of Prussia, PA) diluted in the blocking buffer were incubated with the T cells at 4°C overnight. The slides were them washed 3 times with PBST buffer, each for 5 minutes, followed by the incubation of Alexa Fluor568-conjugated donkey-anti-rabbit IgG (1:100) and Alexa Fluor488-conjugated donkey anti-mouse IgG (1:100,Carlsbad, CA) at room temperature for 60 minutes. The slides were washed 3 times with PBST buffer and mounted with VectorShield mounting media (Vector Laboratories, Newark, CA), and examined with an Olympus fluorescence microscope (Olympus America, Bartlett, TN).

### Live PDA tumor slice preparation

Tumor slices were prepared as previously described (11, 21). Fresh pancreatic tumor tissue harvested from *KPC* or *KPCT* genetically engineered mice was kept on ice in 1x PBS with 10 µg/ml STI for support before slicing using a tissue vibratome (Campden Instruments, London, UK). The harvested tumor is superglued to the vibratome cutting stage with 1.5% agarose gel as support. The stage is filled with ice-cold 1X PBS for tumor slicing, and tumor tissue is sectioned at a 350 µm thickness with a vibratome speed of 0.3 mm/s and frequency of 8mm. Sliced tumors are placed in sterile ice-cold 1X PBS with STI for transport. Tumor slices are then carefully transferred and cultured in a 6-well, or 12-well tissue culture plate with an organotypic tissue culture insert (MilliporeSigma, US) coated with 3 mg/ml of rat tail collagen type 1 neutralized with 1mM HEPE in 2X PBS at a 1:1 ratio and completed with 1X PBS. Multiple slices are placed flat on 0.4 µm, 30 mm diameter cell culture inserts, while a single tumor slice can be placed on 0.4 µm, 12 mm diameter cell culture inserts. Tumor slices were cultured in prewarmed RPMI 1640 supplemented with 10% FBS, 1% penicillin and streptomycin (P/S), 5 µg/mL plasmon (Invivogen), and 10 µg/ml of STI at 37 °C in a 5% CO2 humidified incubator. The tissue culture medium is changed daily, and the pancreatic tumor slices can be cultured for up to 8 days while preserving tissue integrity and cell viability.

### T cell migration assays

To assess migration, T cells were labeled with 2 µM CellTracker™ Green (ThermoFisher Scientific, MA) and seeded into either 3D collagen matrices or PDA tumor slices derived from *KPC* or *KPCT* mice. Collagen matrices were prepared from high-concentration rat tail collagen type I, neutralized with an equal volume of HEPES buffer, and allowed to polymerize before cell seeding. *KPC* tumor slices were stained with 5 µM CellTracker™ Red CMTPX, while KPCT slices required no staining due to endogenous tdTomato expression specifically in carcinoma cells. Following a 1-hour incubation at 37 °C, collagen matrices or tumor slices were rinsed with warm L-15 medium containing 1% FBS to remove non-infiltrating T cells and transferred to 35-mm imaging dishes. A tissue harp was used to stabilize the gels or slices. For live-cell imaging, 5 mL of L-15 medium supplemented with 10% FBS, 1% penicillin/streptomycin, 5 µg/mL Plasmocin, and 10 µg/mL STI was added. The imaging chamber was maintained at 37 °C. T cell migration was recorded at up to two regions of interest per sample, capturing 100 µm z-stacks (5 µm step size) at 1-minute intervals for up to 8 hours.

For live cell imaging, Multiphoton Laser-Scanning Microscopy (MPLSM) to simultaneously generate multiphoton excitation (MPE) of fluorescence (for fluorescent proteins) and second harmonic generation (SHG; for collagen). Imaging was performed on the Provenzano lab custom-built multi-photon laser scanning microscope (Prairie Technologies / Bruker) using a Mai Tai Ti:Sapphire laser (Spectra-Physics) and 4 channel detection as described (11, 30)to simultaneously generate MPE and SHG to visualize cells and collagen, respectively, at an excitation wavelength of 880nm.

### Migration analysis

Briefly, time-lapse images of the T cells migration in 3D collagen or *KPC/KPCT* tumor slices are post-proceeded in Fiji. Stage drifts are corrected using the 3D correction plugin in Fiji using the SHG channel for registration. T cell movements can be tracked using the Trackmate plugin in Fiji, where dynamic threshold and filters can be used to exclude erroneous tracks that might be generated by imaging artifacts or the tracking program(31). The post-processed cell tracking data are then analyzed by fitting to a persistent random walk model using overlapping intervals in the MATLAB model as previously described (32). Briefly, the mean squared displacement (MSD) for cells over time interval it is taken from the average of all squared displacements such that 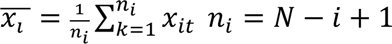, where i is the number of overlapping time intervals of duration *t_i,_* and N is the total number of time intervals. The mathematical representation of the persistent random walk model is the following: 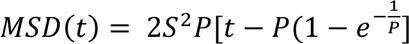, where S is the migration speed, and P is the persistent time. The motility coefficient can be written as such that: 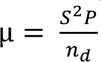, where n_d_ is the dimensionality of the random walk. For this study, the motility, speed, and persistence times for x, y, and z directions were obtained by fitting the model separately into three orthogonal directions; thus, n is one for each case. The total speed of each cell can be calculated by taking the square root of the squared sum of speed in each of the three directions, x, y, and z. The equation can be written as such that: 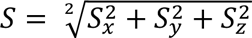. The total motility of each cell can be obtained by adding the motility of each direction and can be written as such that: µ = µ*_x_* + µ*_y_* + µ*_z_*.

### T cell metabolism and membrane tension by multiphoton FLIM

T cells were cultured overnight with or without ImmunoCult human CD3/CD28 T cell activator antibodies (StemCell, Cat. #10971). One million cells of each type were then added to 3mg/mL collagen gels in 24 well plates and allowed to infiltrate for 1h. The collagen matrix was removed from the well, rinsed, and placed in a 35mm dish under a tissue harp in L15 media for imaging. For each gel, 4-6 Z stacks in 5um steps were collected, using MPLSM system described in the T cell migration assays section that is equipped for multiphoton fluorescence lifetime imaging microscopy (FLIM). Excitation at 750nm was used for NAD(P)H excitation and emission was collected through a 460/50 bandpass filter into a GaAsP photon counting PMT (30). From the PMT, photon data were time tagged using SPC-150 photon counting electronics (Becker & Hickl GmBH).

Each lifetime image was imported to SPCImage 8.9 (Becker & Hickl GmBH) and the fluorescence lifetime was deconvolved from the instrument response function using a bin of 16 pixels (4×4) and fit to a two component exponential decay model corresponding to the short and long lifetime components of NAD(P)H. α1 is the fractional contribution of the short lifetime component, was exported as text images, and used as an indicator of T cell metabolism and T cell activation as previously described (25). Using the Fiji package of ImageJ (33),NAD(P)H intensity images were thresholded to include cytoplasm but omit the dim nucleus and background and converted to binary. Binary images were divided by 255 so included pixels equal 1 and excluded pixels equal 0 and then multiplied by the corresponding α1 images. α1 values from all remaining pixels were averaged and the averages compared using an ordinary one-way ANOVA with Tukey’s multiple comparisons test in GraphPad Prism 10.4.1.

### Flow Cytometry to assess T cell differentiation, activation, and exhaustion

Isolated human T cells were stained using standard procedures with the following panel and analyzed by flow cytometry using a BD FACSymphony A3 Cell Analyzer (Becton, Dickinson and Company [BD]) at the University of Minnesota – Twin Cities, University Flow Cytometry Resource: Live/Dead Ghost 780 (Cytek/TONBO, Cat. # 13-0865-T100), CD45-BUV661 (BD, Cat. # 705178), CD3-BUV396 (BD, Cat. # 563548), CD4-APC-R700 (BD, Cat. # 565995), CD8-RB613 (BD, Cat. # 571090), CD45RA-RB705 (BD, Cat. # 757395), CCR7-BV421 (BD, Cat. # 566744), CD27-RY610 (BD, Cat. # 571334), CD28-BV711 (BD, Cat. # 563131), CD62L-BV650 (BD, Cat. # 563808), CD25-PE-Cy7 (BD, Cat. # 560920), CD127-BUV496 (BD, Cat. # 749984), CD103-FITC (BD, Cat. # 567677), PD1-BV605 (BD, Cat. # 563245), LAG3-BV510 (BD, Cat. # 746609), TCF1-AF647 (BD, Cat. # 566693), and CD69-BUV737 (BD, Cat. # 612818). Gating for T cell differentiation was done as previously described (27).

### 2D T cell killing assay

Equal number of tumor cells were plated in each well and the corresponding ratio of T cells to tumor was added to assess T cell killing. The cells were co-cultured for 48 hours before assaying T cell killing efficacy using flow cytometer (BD Biosciences, NJ). T cells and tumor cells were selected based on size and complexity using the FSC/SSC plot. Tumor cells cultured without T cells were used as a baseline and the percent survival for each of the coculture groups was calculated.

### 3D cancer cell – T cell interaction assay

500K AsPC1 cells (ATCC) were stained with 5 µM CellTracker™ Red (ThermoFisher Scientific, MA) and mixed in 3mg/mL of 3D collagen matrices. The cell collagen mixtures were then transferred to a 24 well plate and were allowed to polymerize overnight in 37°C incubator. CD8⁺ T cells labeled with 2 µM CellTracker™ Green were then plated on top of the collagen matrix and incubated allowed to infiltrate into the matrices for 1 hour. Any T cells remaining outside of the collagen matric after incubation was washed off. T cell, AsPC cell, and collagen were virtualized in using MPE and SHG and imaged for 90 minutes with a 100 µm z section, 1 minute interval. T cell – carcinoma interaction frequency was then analyzed using PhenoTrack, a component of the Provenzano lab TME-CART algorithm suite, that uses T cell morphology, velocity, and physical proximity with carcinoma cells to identify the phenotypic behavior of cells. Taking inputs of T cell tracks and T cell morphology over time from TrackMate in FIJI, PhenoTrack examines T cell velocity, circularity, the change in cell circularity, and the distance to cancer cells at each frame for each T cell. Based on tunable thresholds for each parameter, PhenoTrack assigns a plausible status (migration, stalling, interaction) to each cell at each time point.

### *In vivo* tumor assays

3×10^5^ AsPC1 cancer cells were mixed with 6×10^6^ human cancer associated fibroblasts (CAFs) in 100 µL of 3mg/mL of collagen gel and orthotopically injected into each NSG mouse pancreas on Day 0. Tumor growth was monitored using the M5 compact MRI (Aspect Imaging) system and mice were enrolled in intervention studies when the tumor reached **XXXX**. Tumor volume (V) was calculated using the formula: V = ½ * L *W^2, where L is the longest diameter and W is the shortest diameter. Mice were randomly assigned into three groups: no T cell control, mesoCAR treatment, and mesoCAR RhoA(Q63L) treatment. When tumor reached enrollment criteria, 4×10^6 mesoCAR or mesoCAR RhoA(Q63L) T cells were intraperitoneally injected. When control tumros reached endpoint criteria, mice were sacrificed, and tumor were harvested to analyze T cell infiltration using flow cytometry and immunofluorescent staining.

For quantifying T cell infiltration number, a small piece of tumor tissue was digested in tumor dissociation solution (Miltenyi Biotec) for 30 minutes at 37°C and passed through a 70µm cell strainer. Isolated cells were stained for live/dead and anti-CD34 CAR T marker. Remaining tumor tissue were formalin fixed and parafilm embedded (FFPE) for histology analysis. FFPE sections were stained for pan-cytokeratin-FITC (1:200; Sigma, F3418) and human CD34-PE for RQR8 (1:100; Thermo Fisher, MA1-10205), and whole slide fluorescent imaging were taken using Axio Scan System (Zeiss). Images were post-processed in FIJI to remove background and quantify RQR8 positive cells.

### Statistical analysis

One-way ANOVA followed by Tukey post hoc analysis is used to determine if there is a statistical difference between multiple groups with log transformation as appropriate, and unpaired two-way Welch’s t-test or Student’s t-test, based on population variance, is used to analyze pairwise comparisons. All statistical analyses presented here were completed using Prism 10 (GraphPad Software, Inc) software. The difference is considered significant if p < 0.05.

## ACKNOWLEDEMENT

This work was supported by the NIH P01CA254849 Project 2 (PPP), U54CA268069 CCBIR Center for Multiparametric Imaging of Tumor Immune Microenvironments (C-MITIE), R01CA286615 (PPP), and American Cancer Society and the American Cancer Society – CIO’s Against Cancer MN Chapter - Discovery Boost Grant, DBG-24-1322329-01-IBCD (PPP). We thank a Brianna teDuits, Paarth Dodhiawala, and Priyanila Magesh for assistance with mouse surgeries.

## SUPPLEMENTARY INFORMATION

**Supplementary Figure 1:**
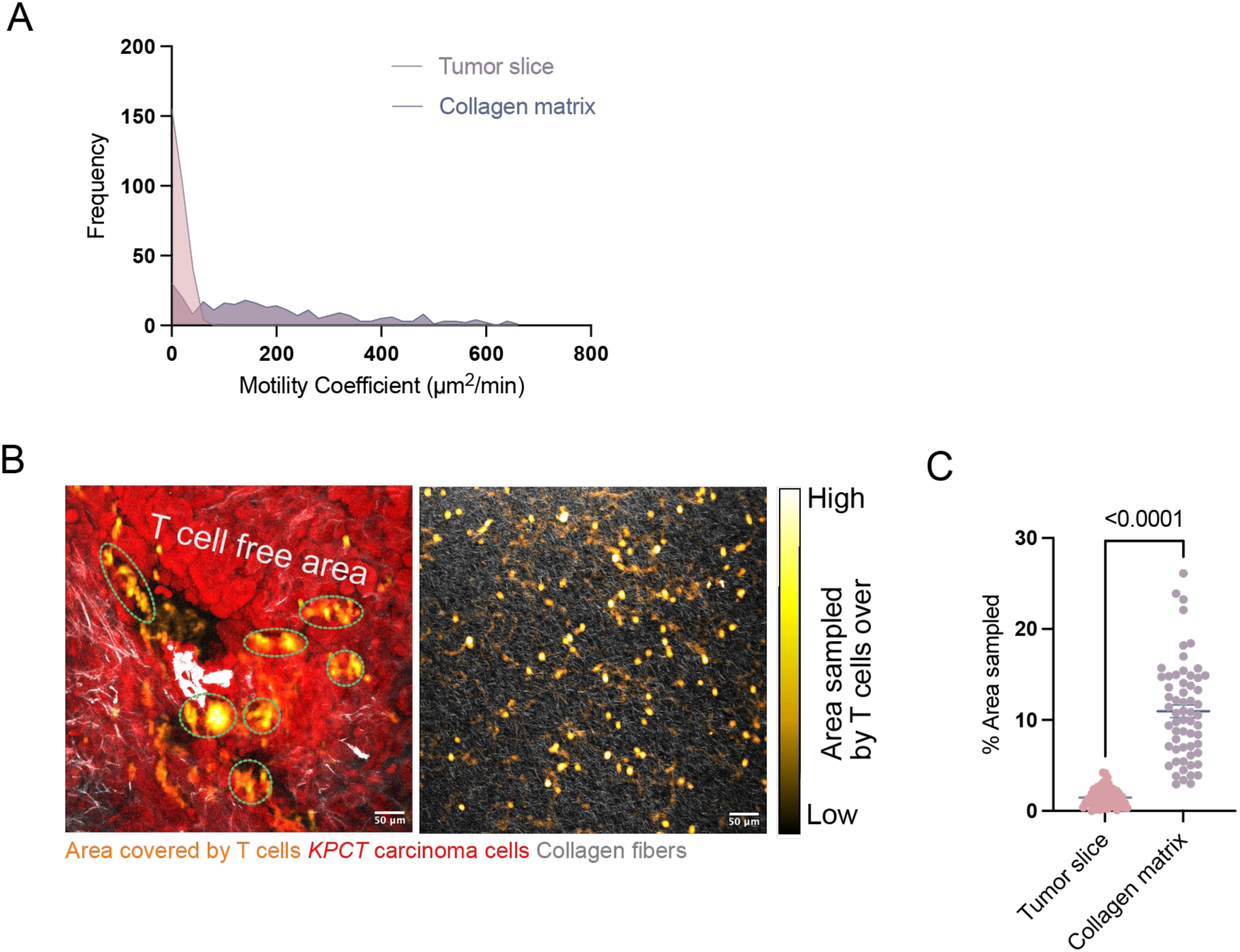
Wild-type CD8+ T cell migration is limited in PDA TMEs. (**A**) Distribution of wild-type murine CD8+ T cell motility coefficient in tumor slice (pink) and 3D collagen matrices (purple). (**B**) Representative images showing spatial sampling by T cells in live mouse PDA tumor slice (1 hours) and in 3D collagen matrix (1 hour). Brighter color indicates higher T cell presence. (**C**) Quantification of total area covered per field of view overtime by CD8+ T cells in tumor slice (Left, 1 hour) and in 3D collagen matrix (right, 1 hour). Data are presented as mean ± s.e.m. Statistical comparisons were performed using unpaired two-tailed t-tests with Welch’s correction.

**Supplementary Figure 2:**
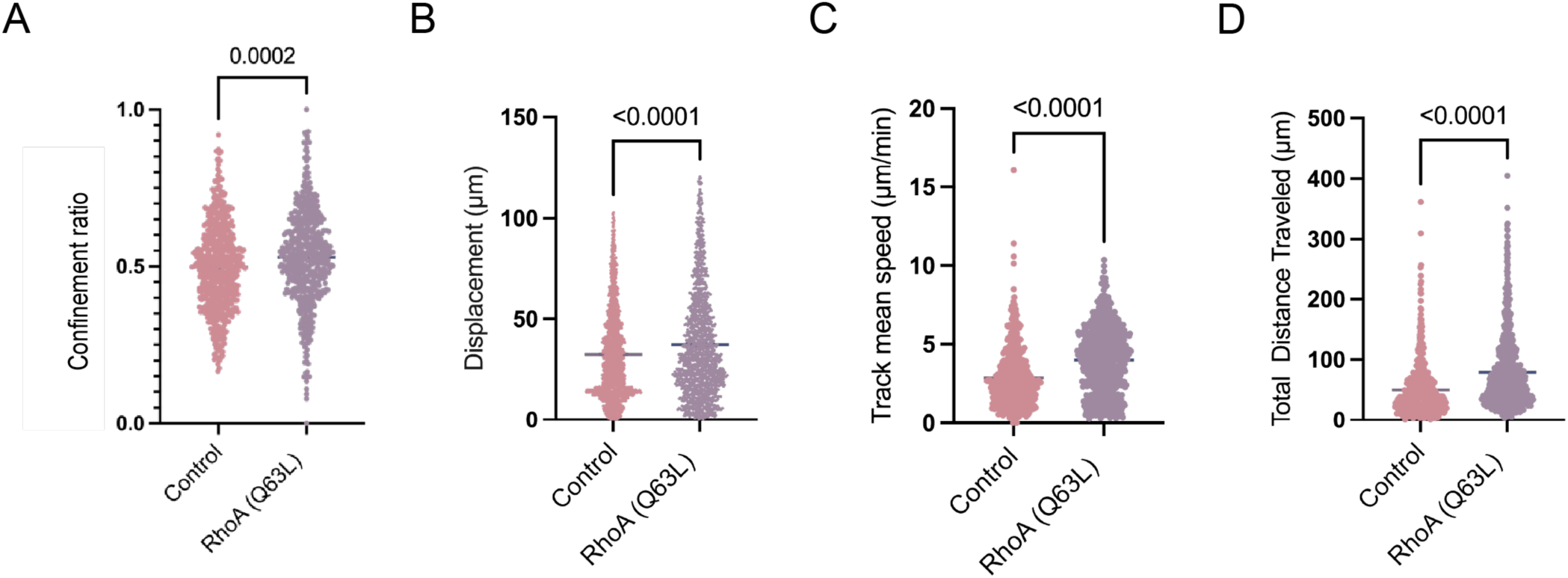
Human RhoA(Q63L) modified T cells are more migratory. **(A)** Expression of constitutively active Rho results in (**A**) **less confined migration,** (**B**) increased displacement, (**C**) mean speed, and (**D**) total distance traveled (**D**). Data are presented as mean ± s.e.m.; statistical significance was determined using unpaired two-tailed t-tests with Welch’s correction.

**Supplementary Figure 3:**
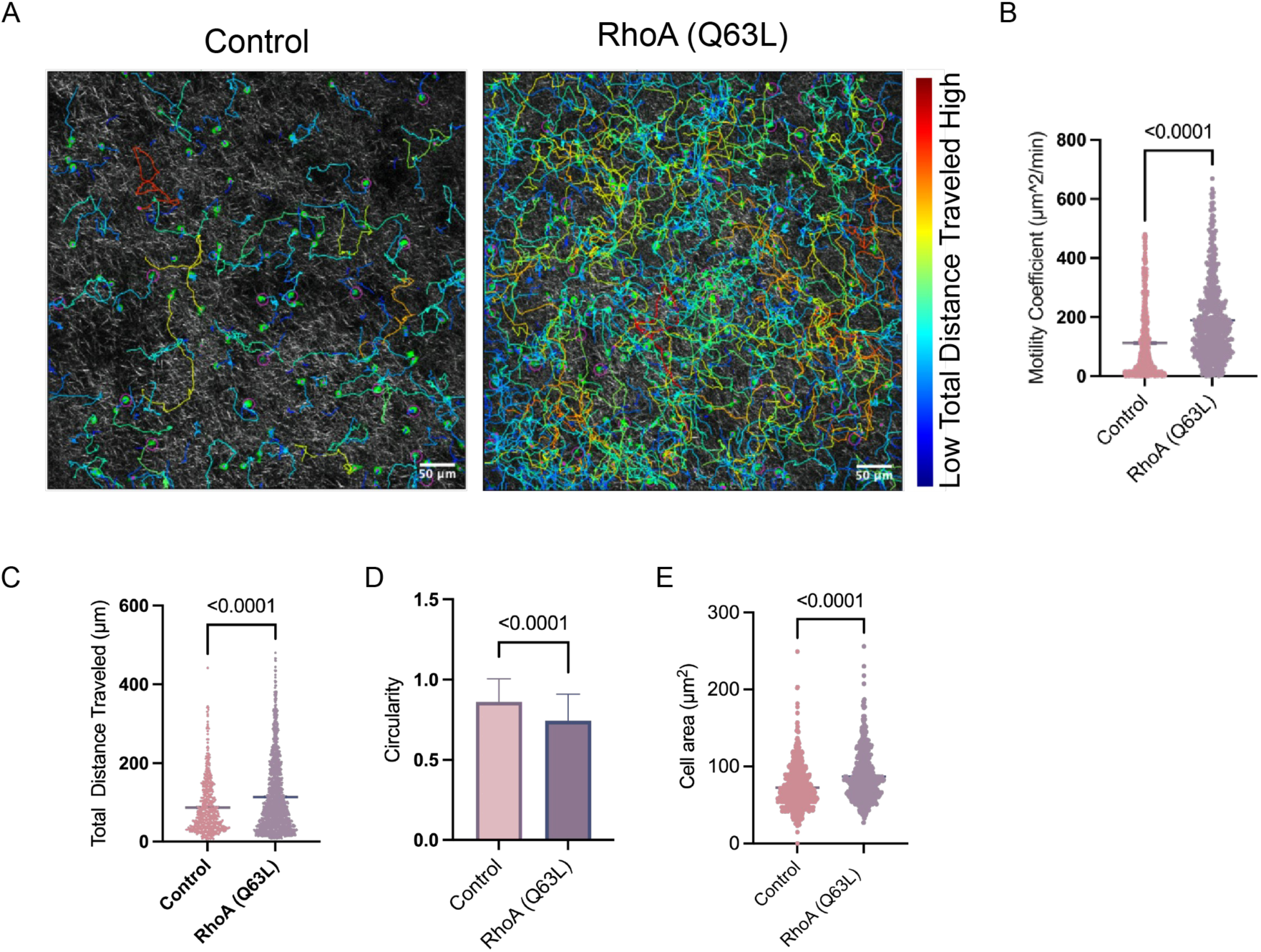
Mouse CD8+ T cells expression constitutively active Rho have higher motility in 3D matrices. (**A**) Representative multiphoton time-lapse images of murine RhoA(Q63L) and control CD8⁺ T cells migrating in 3D collagen matrices. Cell trajectories are overlaid and color-coded by total distance traveled (blue = short; red =long). Images were acquired every 1 min for 1 h. Colors: green, T cells; gray, collagen fibers. (**B**) Motility coefficient of murine RhoA(Q63L) and control CD8⁺ T cells migrating in 3D collagen matrices. Each dot represents an individual tracked T cell. (**C**) Total distance traveled by murine RhoA(Q63L) and control CD8⁺ T cells in 3D collagen matrices. Each dot represents an individual tracked T cell. (**D**) Average circularity of murine RhoA(Q63L) and control CD8⁺ T cells in 3D collagen matrices during the imaged period. A value of 1.0 indicates a perfect circle, and the value approaches 0 for elongated cells. (**E**) Cell size of murine RhoA(Q63L) and control CD8⁺ T cells in 3D collagen matrices measure using particle analyzer in FĲI. Data are presented as mean ± s.e.m.; statistical significance was determined using unpaired two-tailed t-tests with Welch’s correction.

**Supplementary Figure 4:**
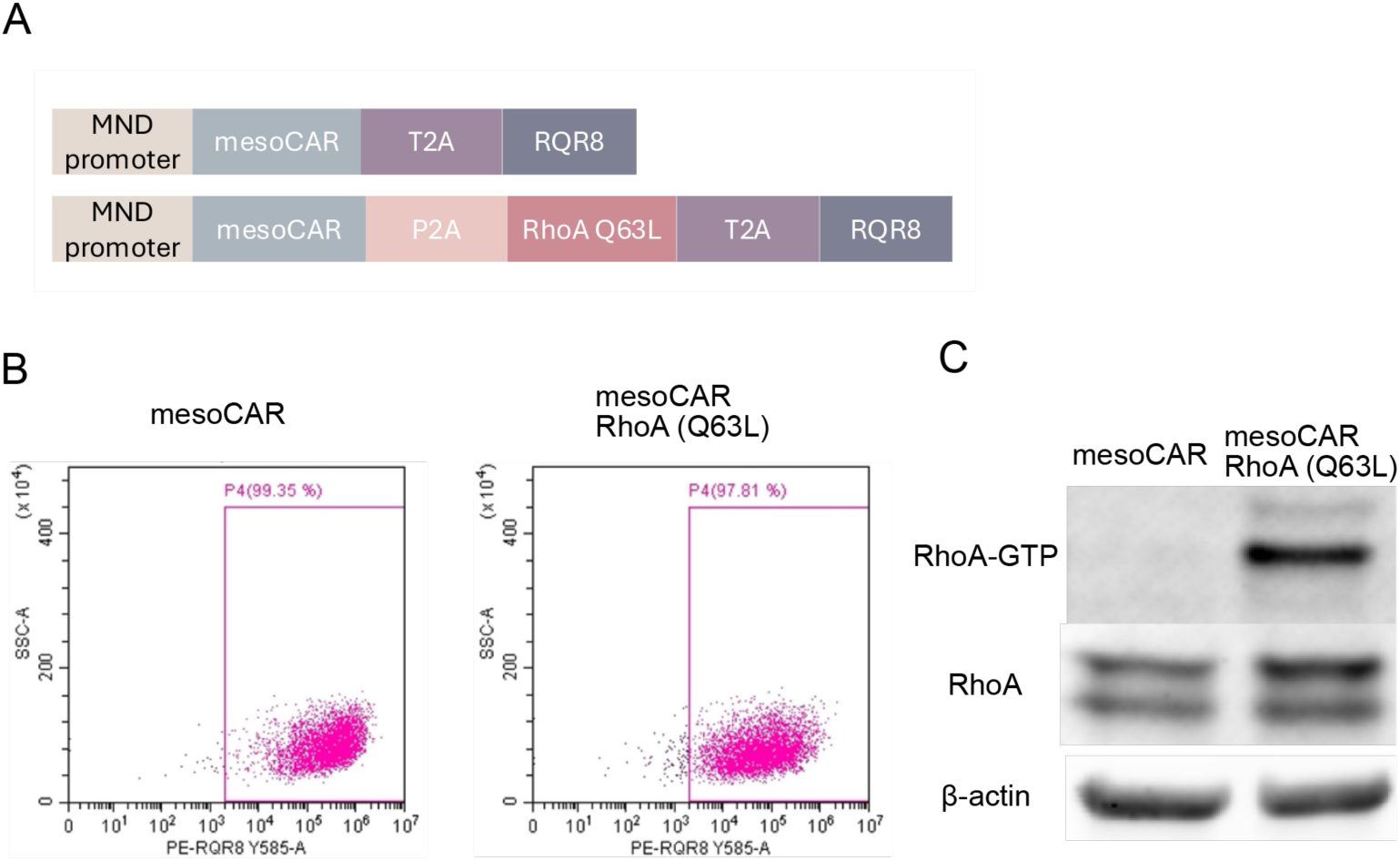
Characterization of mesoCAR and mesoCAR RhoA(Q63L) T cells by flow cytometry and RhoA activation assays. (**A**) Schematic representation of vector constructs encoding mesothelin-targeted CAR (mesoCAR) T cells with or without the RhoA(Q63L) mutation. (**B**)Representative flow cytometry data of purified mesoCAR RQR8+ T cells using PE -conjugated anti-CD34 (QBend10) antibody. (**C**) Western blots of whole cell lysate and RhoA-GTP pulldown samples with RBD beads with antibodies of RhoA and β-actin.

**Supplementary Figure 5:**
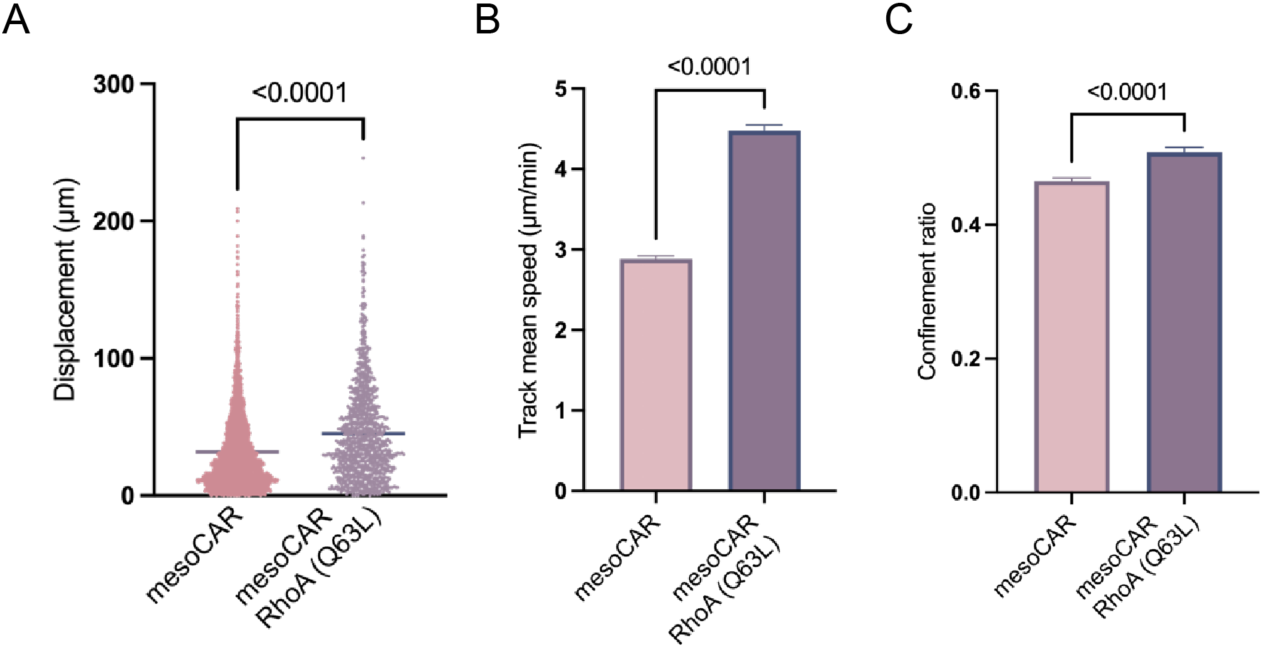
Increased motility of human mesoCAR RhoA(Q63L) CAR T cells in 3D collagen matrices. (**A**) The mesoCAR RhoA(Q63L) cells showed increased displacement, (**B**) mean speed, and (**C**) less confined movement.

**Supplementary Figure 6:**
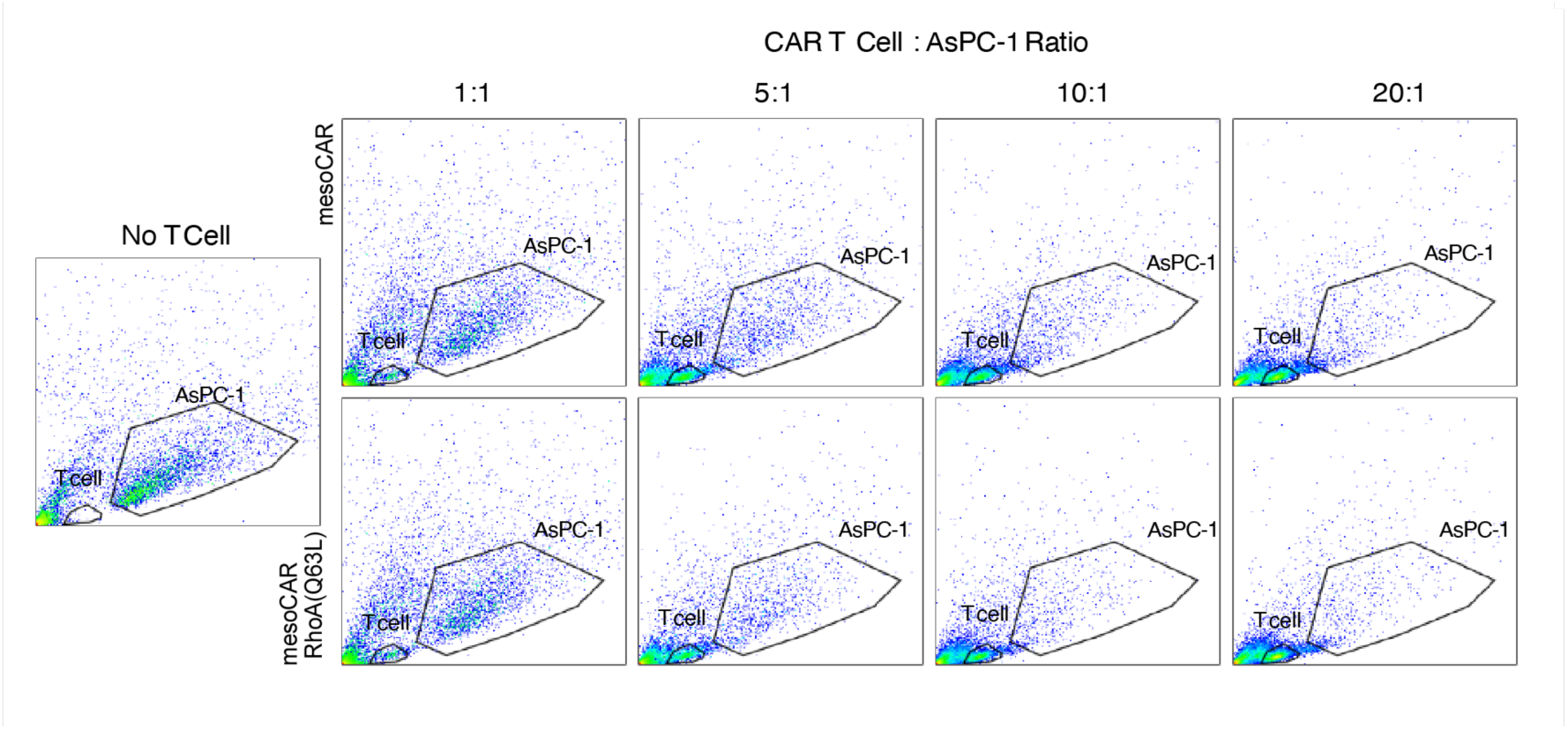
2D cytotoxicity. MesoCAR and mesoCAR RhoA(Q63L) T cells were co-cultured with AsPC-1 at varying effector-to-target ratios. The percentage of viable AsPC-1 cells is shown relative to no-T-cell controls.

### The Smoluchowski limit of immune cell mediated killing

To execute their antitumoral and infection-clearing function, cytotoxic T cells need to come into direct physical contact with cancer/infected cells. Thus, a key step in the immune response to tumors (and infections) is the physical migration of T cells through tissues containing cancer/infected cells to locate, contact, and kill cancer/infected cells. In the case where the target cells are sparsely distributed (diffusely infiltrated) in the tissue, the process of simply finding target cells can potentially become onerous and extremely time consuming, thus permitting cancer cells more time to proliferate and migrate. At some point the T cells will not be able to contain the tumor and spreading will proceed beyond the immune system’s ability to control it. Determining the factors that limit the rate of collision between T cells and cancer cells is important to optimizing T cell function in diffusely infiltrative tumors.

The immune response can be described as a process of collision between two (approximately) diffusing particles, the cancer cell and the T cell. Assuming that these particles can be approximated as spheres whose migration is modeled as a random walk, we can apply diffusion theory to estimate the average time that a T cell spends looking for (hunting) a cancer cell before contact. The association rate constant, k_on_ (units:M^-1^ s^-1^) for diffusing spheres is given by (Smoluchowski, Z. Phys. Chem., 1917; Schlosshauer and Baker, Prot Sci, 2004)

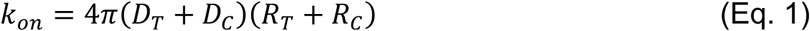

where D_T_ and D_C_ are the random motility coefficients of the T cells and cancer cells (units m^2^/s), respectively, and R_T_ and R_C_ are their radii. Given our estimates of the random motility coefficients and radii of the rhoA-WT T cells and cancer cells in our experiments, we estimate

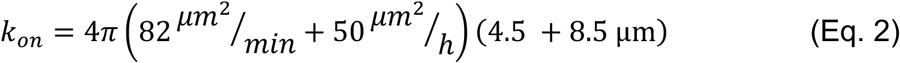

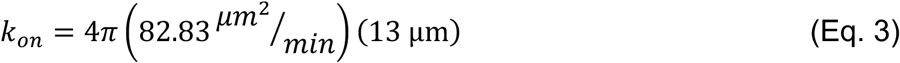

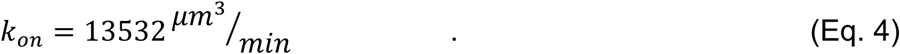

From this analysis, it is evident that the association rate constant is controlled by the T cells rather than the cancer cells, since the migration speed of the former is more than an order of magnitude greater than that of the latter.

The concentration of cancer cells in our experiments is estimated to be

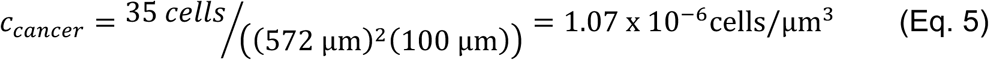

where the spatial dimensions are given by the spatial dimensions of the volume containing 35 cancer cells given by the xy dimensions of the field (572 µm)^2^ and the z depth of the z-stack (100 µm). Thus, the pseudo-first order rate constant for meso CAR T cells contacting cancer cells at this cancer cell concentration is given by

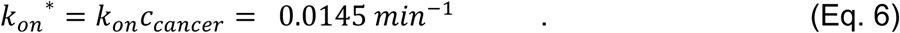

Equivalently, it can be stated that the average time that a T cell migrates before contacting a cancer cell, <τ>, under these conditions is given by

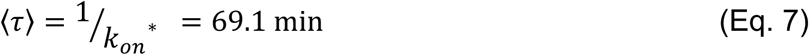

meaning that T cells spend 69.1 min, on average, searching for a cancer cell to kill before they actually contact one. Given that the average dwell time of a mesoCAR T cell in contact with a cancer cell is 9.3 min, the average fraction of the time that an average mesoCAR T cell would spend searching is given by

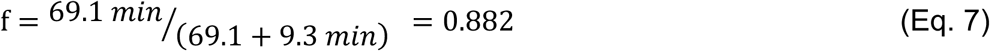

so that we estimate mesoCAR T cells spend about 88% of their time searching for cancer cells and only 12% of it in contact, hopefully killing the cancer cell (assuming that contact events always lead to killing, which may be optimistic). This estimate means that T cells under normal conditions may very well be limited in effectiveness in responding to solid tumors by the shear amount of time they spend searching for them, implying that greater T cell motility, as we observed for mesoCAR Rho(Q63L) T cells would make the T cells more therapeutically effective.

Repeating the above calculations for mesoCAR Rho(Q63L) T cells, which we found were faster than mesoCAR T cells (139 µm^2^/min vs 82 µm^2^/.min), we find the pseudo-first order rate constant for mesoCAR Rho(Q63L) T cells contacting cancer cells at this cancer cell concentration is given by

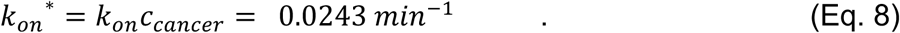

Correspondingly, the average time that a mesoCAR Rho(Q63L) T cell migrates before contacting a cancer cell is given by

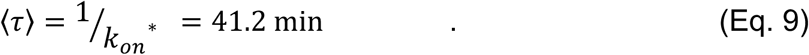

meaning that mesoCAR Rho(Q63L) T cells spend about 41.2 min, on average, searching for a cancer cell to kill before it contacts one. Given that the average dwell time of a mesoCAR Rho(Q63L) T cell in contact with a cancer cell is 21.5 min, the average fraction of the time that a mesoCAR Rho(Q63L) T cell would spend searching is given by

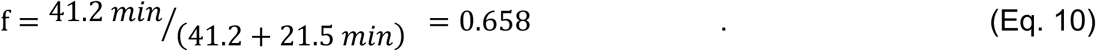

so that we estimate mesoCAR Rho(Q63L) T cells spend about 66% of their time searching for cancer cells and 34% in contact with them. This estimate means that mesoCAR Rho(Q63L) T cells are expected to be much more effective in responding to sparsely distributed cancer cells, by virtue of the reduced amount of time they spend searching for them, compared to mesoCAR T cells.

